# Sex-dependent effects of stress on insular cortex-to-nucleus accumbens synaptic plasticity

**DOI:** 10.1101/2023.08.28.555067

**Authors:** Manon Gauthier, Emilie Dugast, Virginie Lardeux, Kevin Letort, Laure Belnoue, Eric Balado, Marcello Solinas, Pauline Belujon

## Abstract

Stress is an important risk factor for the development of psychiatric disorders and men and women tend to react differently to stress. Sex differences are also observed in many stress-related psychiatric disorders such as depression, anxiety disorders or addiction. Therefore, identifying specific neuroadaptations induced by stress, in males and females, is a necessary step to the understanding of stress-related sex dimorphism in these disorders. Here, we tested the hypotheses that acute stress could affect plasticity in the anterior insular cortex (aIC)-nucleus accumbens core (NAcC) pathway, two structures involved in the stress response, in a sex-dependent manner. Using *in vivo* extracellular recordings in anesthetized rats, we show that synaptic plasticity in the aIC-NAcC pathway is different between male and female rats. Whereas in males, long-term potentiation and long-term depression were equally induced, in females, there was mostly a long-term potentiation induced. Moreover, stress affected synaptic plasticity in the aIC-NAcC differently in male and female rats. In males, stress induced a loss of long-term-depression that lasted for at least 24h, whereas in females, stress induced less neurons displaying LTP, which did not last.

These results demonstrate that integration of aIC information to NAcC is different between males and females. This study provides mechanistic support for differential reactivity to stress between males and females that may relate to stress-related psychiatric disorders and sex dimorphism in these disorders.

## INTRODUCTION

Stress response involves a neuro-behavioral cascade which is elicited when the organism is confronted with a perceived harmful situation. This comprises a range of neural and endocrine changes including activation of the autonomic nervous system and the hypothalamic-pituitary-adrenocortical (HPA) axis (Chrousos and Gold, 1992) and selective attention (Chajut and Algom, 2003). This stress response activates appropriate avoidance, coping and restorative processes that will remain until homeostasis is restored. Although acute stress does not usually constitute a danger to an organism’s health, persistence or inappropriate activation of the stress system may represent one of the risk factors for mental disorders (Smith and Pollak, 2020). Because of the very high health cost of psychiatric disorders for individuals and for the society, it is necessary to identify the neurobiological mechanisms that render individuals vulnerable to stressors.

Clinical and preclinical research has demonstrated significant sex differences in the response to stress, and sexual dimorphism in stress-related disorders (Bangasser and Valentino, 2014). For example, sex differences have been observed in the expression, distribution, signaling and trafficking of corticotropin-releasing factor (CRF, which activates the HPA axis) receptor (Bangasser and Wiersielis, 2018).

Moreover, previous studies have proposed that stress alters the central representation of bodily processes (Kessler et al., 2012). The perception of these internal body processes, integrated in the insular cortex (Craig, 2009) plays an important role in mental disorders such as depression, anxiety or addiction (Khoury et al., 2018). Interestingly, differences in interoceptive accuracy has been shown between men and women (Prentice et al., 2022). Compared to males, females exhibit worse interoceptive accuracy but show greater emotional self-awareness (Thompson and Voyer, 2014).

The insular cortex is involved in interoceptive stimuli processing and the anterior insular cortex (aIC) in particular is a cerebral hub encoding interoceptive information to generate emotional awareness (Craig, 2009; Critchley et al., 2004; Gu et al., 2013). Neuroimaging studies in humans have highlighted the IC as a core region affected across many stress-related psychiatric disorders including addiction (Naqvi et al., 2014), depression (Sliz and Hayley, 2012) or anxiety disorders (Terasawa et al., 2013). Recently, Gehrlach and collaborators demonstrated that the anterior insular cortex (aIC) sends dense projections to the core of the NAc (NAcC) (Gehrlach et al., 2020), a limbic-motor hub, where learned associations of motivational significance are converted into goal-directed behavior (Mogenson et al., 1980). The NAc receives convergent inputs from different structures involved in contextual, emotional and cognitive processes, which can be altered by stress (Floresco, 2015). Acute stress has been shown to increase dopamine release in the NAc (Stevenson and Gratton, 2003) and alter synaptic plasticity and gating of different glutamatergic inputs in the NAc (Gill and Grace, 2013), suggesting stress-induced alterations that could alter context-dependent behaviors.

Considering stress-dependent sexual dimorphism, we hypothesized that the integration of information from the aIC in the NAc would be different between males and females. For this, we investigated changes in synaptic plasticity in the aIC-NAcC pathway after restraint stress.

### Experimental procedures

#### Animals

Adult (> 9 weeks) male and female Sprague-Dawley rats (Janvier, France) arrived at least 4 days before experimentations. They were housed two per cage from arrival to the beginning of the stress protocol under a 12h-light/-dark cycle (on at 7:30 am) with controlled temperature (22°C) and humidity (47%). Rats had *ad libitum* access to food and water in their homecage. All experiments were conducted during the light phase and in accordance with the European Union directives (2010/63/EU) for the care of laboratory animals. All experimental protocols were approved by the local ethics committee (COMETHEA).

#### Stress protocol

Rats were subjected to a restraint stress which consisted of a 2h-restraint session in a cylindric plexiglas size-adjusted tube (6cm diameter for females and 7cm for males, Phymep, Paris, France). For electrophysiology, three groups were distinguished: a restraint stress group (RS) where experiments were performed immediately after restraint, restraint stress + 24h group (RS24) where experiments were performed 24 hours after restraint and a control group which remained in their home-cage.

#### cFos experiments

Rats were habituated to the experimental room in their homecage for 1h. Rats were then restrained for 2h and replaced 1h in their homecage in the same experimental room. A Control group underwent the same protocol without being restrained.

#### Tracer injection

Rats received a subcutaneous injection of the nonsteroidal anti-inflammatory ketoprofen (2.5mg/kg, s.c.). and were anesthetized with isoflurane (5% induction, 2.5% maintenance) in O_2_ before being placed in a stereotaxic frame. Animals were placed on a heating pad for the entire procedure. Lidocaine (1mg/kg) was injected locally before incision of the skin. Then, craniotomies were performed above the brain areas of interest. A retrograde tracer cholera toxin subunit B (CTB, 0.5 μl, List Labs, Campbell, CA, USA) was injected into the right NAcC [anteroposterior (AP), +1.1mm for males and +1.2mm for females from bregma; lateral (L), 2.2mm from the midline; dorsal (D), -7.4mm from the skull surface]. Following injection, the incision was sutured, and the rats were placed in their homecage for 2 weeks before brain collection.

#### Immunohistochemistry

Rats were isoflurane-anesthetized and were euthanized with a lethal dose of pentobarbital (>150 mg/kg, i.p.). Rats were perfused intracardially with NaCl 0.9% and paraformaldehyde 4%. Brains were extracted and the tissue was left 24 hours in paraformaldehyde 4% and cryoprotected 48h in a 25% sucrose solution. Once saturated, brains were coronally sliced (40μm) using a microtome (HM 450, Thermo Fischer Scientific, Illkirch-Graffenstaden, France). Brain slices were washed three times in phosphate buffered saline (PBS 1X).

For cFos immunostaining: a first incubation of 30min in H_2_O_2_ 1.2% + TrisHCl 0.01M pH7.6 was performed to inactivate peroxidases. Three washes in PBS 1X were done before a second incubation with bovine serum albumin (BSA) blocking buffer solution (PBS 1X-Triton 100X 0.3% + BSA 3%) during 1h30. Brain slices were incubated overnight under agitation at 4°C in a solution of primary antibody mouse anti-cFos (1/10000 diluted in BSA buffer solution, C-10 sc271243, Santa Cruz Biotechnology, Dallas, TX, USA). After three washes, secondary antibody incubation was done with a donkey anti-mouse biotinylated antibody (1/1000 diluted in BSA buffer solution, 715-065-150, Jackson, Cambridge, UK) under agitation for 1h30 at room temperature. Sections were washed and incubated 1h with avidin-biotin complex (ABC kit, Vector, PK-6100, Vector laboratories, Newark, CA, USA). Following PBS 1X washes, sections were incubated for 30min in 3,3’-diaminobenzidine-4HCl (DAB substrate kit, SK-4100, Vector laboratories, Newark, CA, USA) containing H_2_O_2_ in Tris Buffer, and three washes in PBS. Brain slices were mounted on gelatinized slides and coverslipped with depex (Sigma-Aldrich, Saint-Quentin-Fallavier, France). Slides were imaged by a transmitted light microscope (Zeiss Imager.M2, Zeiss, Paris, France).

For CTB and NeuN immunostaining: a first incubation with bovine serum albumin bocking buffer solution (PBS 1X-Triton 100X 0.3% + BSA 3%) during 1h30. Braincuts were incubated overnight under agitation at 4°C in a solution of primary antibodies: goat anti-CTB (1/2000 diluted in BSA buffer solution, 703, ListLabs, Campbell, CA, USA) and rabbit anti-NeuN (1/500, ab177487, abcam, Paris, France). After three washes, secondary antibodies incubation was done with a solution of donkey anti-goat antibody (Alexa fluor 568, 1/1000 diluted in BSA buffer solution, A11057, Invitrogen, Waltham, MA, USA) and donkey anti-rabbit antibody (Alexa fluor 647, 1/500, A31573, Invitrogen, Waltham, MA, USA) under agitation for 1h30 at room temperature. Sections were washed and mounted on SuperFrost plus slides and coverslipped with depex (sigma, Saint-Quentin-Fallavier, France). Slides were imaged by a fluorescent microscope, Slideview Olympus VS200 microscope (x10 tiles).

#### Quantification of staining

To quantify cFos labelling, we acquired 3 slices (right and left hemispheres) per rat containing the paraventricular nucleus of the thalamus (PVN). Quantification was done with ImageJ (Schneider et al., 2012) and the values obtained were averaged per rat.

To quantify retrograde labelling and to determine the percentage of neurons positive to CTB, we acquired 3 aIC-slices of CTB and NeuN labelling (ipsilateral to the injection site) per rat, with a total of 5 males and 5 female rats. We quantified CTB- and NeuN-positive cells separately with QuPath (Bankhead et al., 2017) and an average was calculated per rats.

#### Elevated plus maze (EPM)

The EPM apparatus consisted of four elevated perpendicular Plexiglas arms: two opposite open arms and two opposite closed arms of 50 cm length and 11cm width. All surfaces were black and the wall for the closed arm was 43 cm height. Open arms were around 25 lux illumination. Animals were habituated 30 min in the room where the elevated plus maze (EPM) apparatus was placed. Rats of AR group were habituated to the room, then stressed and placed in the center of the EPM, facing one open arm and then allowed to start exploring the maze freely during 5 min test. Video recordings were analyzed using video tracking software (Viewpoint, Lyon, France) to measure distance and time spent immobile. A manual analysis was performed to measure number of entries and the time spent in open arms (to calculate the percentage) realized by the animal.

#### In-vivo single-unit extracellular recordings

Rats were isoflurane-anesthetized (5% induction, 2.5% maintenance) and placed in a stereotaxic apparatus. Extracellular recording electrodes were pulled from glass micropipettes (capillaries 30-0057 GC150F-10, Harvard Apparatus, Holliston, MA, France; impedance, 8.5-14MΩ) and filled with a 2% Chicago Sky Blue dye (Sigma-Aldrich, Saint-Quentin-Fallavier, France) solution in 2M NaCl. Evoked activity was recorded from the right NAcC (AP, + 1.1-1.4 mm relative to bregma; L, 2-2.3 mm from the midline; D, -6.8-8 mm from the skull surface) according to the brain atlas (Paxinos and Watson, 2017). Signals were amplified 50 times, filtered (low pass: 300 Hz; high pass: 16 kHz; Multiclamp 700b, Axon Instruments, Union City, CA, USA) and digitized at 16 kHz (CED 1401, Cambridge Electronics Design, Cambridge, UK), acquired on a computer using Signal 6.05 software (Cambridge Electronics Design).

#### Stimulation protocol

Concentric bipolar stimulating electrodes (FHC, USA) were lowered in the right aIC (AP, 10° angle, + 3.4 mm from bregma; L, 4.0 mm from the midline; D, 5.0 mm from the brain surface). Stimulation of the aIC was delivered every 2 seconds (0.5 Hz, 300 µs width pulse), while recording electrode was lowered in the NAcC. Only NAcC neurons that responded monosynaptically (latency <30 msec) were used for analysis. aIC-evoked-spike probability in the NAcC was calculated by dividing the number of action potentials by the number of stimulations. The current intensity was adjusted to obtain ∼50 % of evoked response. Baseline aIC-evoked activity was recorded from NAcC neurons to have at least 10 min of stable recording. Spike probability was calculated at 5-min intervals. After a stable baseline, high frequency stimulation known to induce synaptic plasticity in the cortex-accumbens pathway (HFS, 50 Hz, 2 sec, 300 µs pulse width, Goto and Grace, 2005) was applied to the aIC with a current adjusted to obtain ∼100% of evoked-spike probability. Evoked-spike probability after HFS was recorded for 40 min with the same current used during baseline. Only recordings of 20 min or more post-HFS were analyzed. Changes of +/- 20 % in spike probability for more than 20 minutes were considered changes of synaptic plasticity (Floresco et al., 2001).

#### Histology

Location of the recording electrode was verified via Chicago Sky Blue dye (Sigma Aldrich) electrophoretic ejection (-19 µA constant current, 20 min). Rats were then euthanized with a lethal dose of pentobarbital (>150 mg/kg, i.p.) and brains were collected. The tissue was fixed for 48 hours in 8% paraformaldehyde and then cryoprotected in a 25% sucrose solution. Once saturated, brains were frozen with isopentane, and coronally sliced (30 μm) using a cryostat (CryoStar NX70, Thermo Fisher Scientific). Slices were mounted onto gelatin-coated slides and stained with a solution of neutral red and cresyl violet. Stimulating electrode sites were verified and only neurons recorded in the NAcC and stimulating electrode in the aIC were included in the data analysis (supplementary figure 1).

Estrus cycle of females was determined in order to study whether the female estrus phase influences the experimental results. Cells of vaginal canal were collected using a cotton-tip soaked of distilled water, spread on slides, and stained with crystal violet solution (Marcondes et al., 2002; McLean et al., 2012) (supplementary figure 2).

#### Statistical analyses

Results are presented as mean ± standard error of the mean (S.E.M.). Statistical outliers were identified with Grubbs’s test and excluded from analysis. Normality was checked with the Shapiro-Wilk test and for samples that do not follow a Gaussian distribution, non-parametric statistics were applied with Mann-Whitney, Kruskal-Wallis tests and repeated measures Kruskal-Wallis followed by Dunn’s post-hoc test. When data were normally distributed, they were analyzed by unpaired two-tailed t-tests, one-way, and two-way repeated measures ANOVA (or mixed model when measures were missing) followed by Holm-Sidak post-hoc test. Proportions were analyzed using a chi-square. The significance was set at p < 0.05 and data were analyzed with GraphPad Prism 9.

## RESULTS

### Acute restraint stress increases cFos expression in the paraventricular nucleus of the hypothalamus in male and female rats

Acute restraint stress in rodents has been shown to increase cFos protein expression density in the paraventricular nucleus of the hypothalamus (PVN), which is part of the hypothalamo-pituitary-adrenal axis (HPA) (Zavala et al. 2011). To validate the restraint model used in our study, we measured cFos expression in male and female rats in the PVN after exposure to acute stress.

Male (figure 1B *left*) and female (figure 1B *right*) rats exposed to restraint stress (RS) showed a significant increase in cFos density in the PVN compared to naive rats (Unpaired t-test; males: t=8.797, p<0.05; females: t=4.624, p<0.05). This suggests that the procedure used in the present study activates the stress system in both sexes.

**Figure 1:**
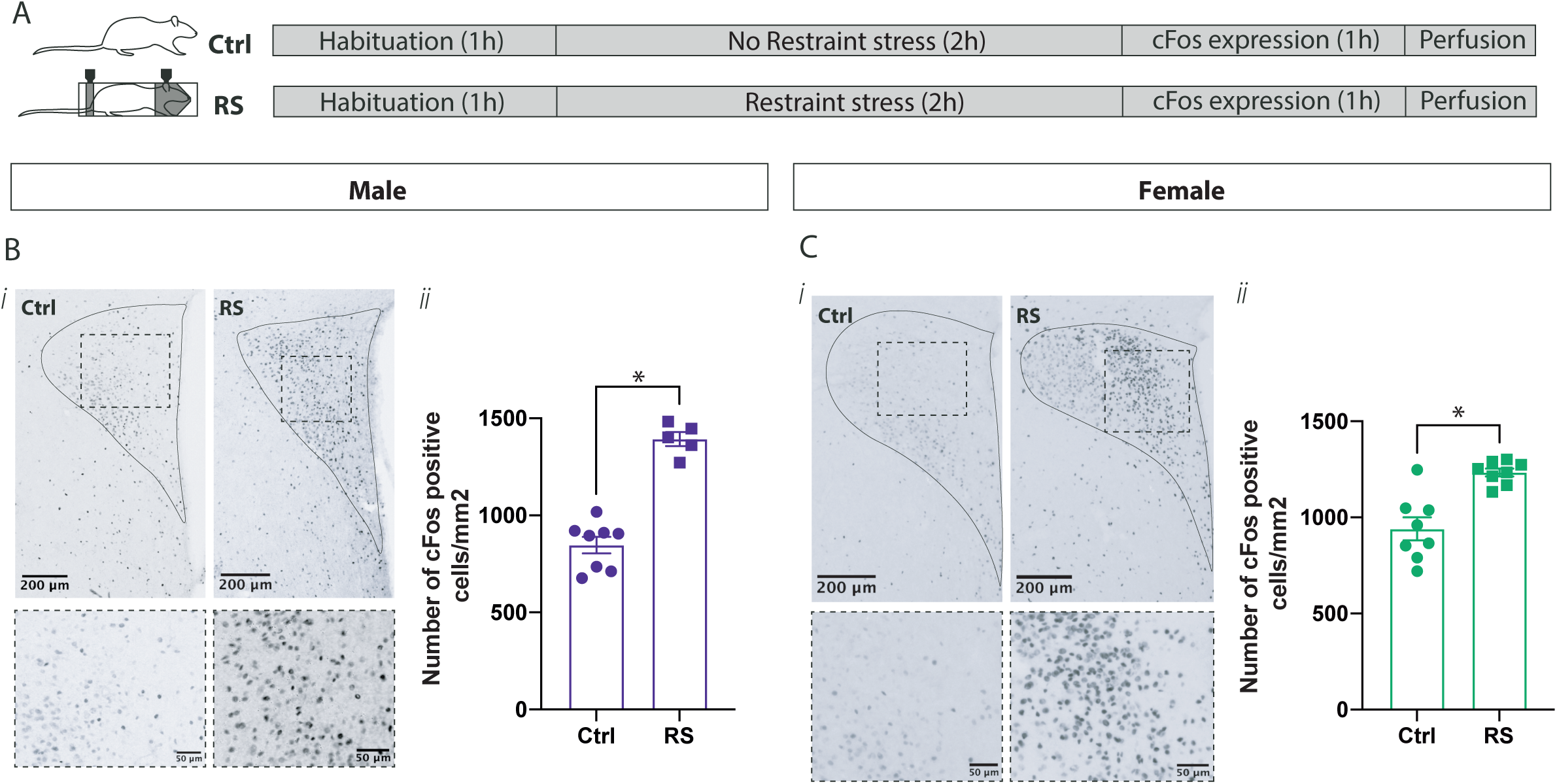
Effect of restraint stress on cFos density in the paraventricular nucleus of the hypothalamus. **(A)** Experimental groups and procedures for quantification of cFos expression in control (Ctrl) and restraint stressed (RS) rats. **(i)** Representative transmitted light images of coronal slices of the paraventricular nucleus of the hypothalamus (PVN) showing cFos expression in a Ctrl and RS rat. **ii)** Impact of RS on mean ± SEM cFos density in the PVN in males (B) and in females (C). Male Ctrl, n=8; Male RS, n=5; Female Ctrl, n= 8; Female RS, n=8; *p<0.05

### Restraint stress decreases exploratory behavior in male and female rats for at least 24h

Restraint stress has been described to increase avoidance of the open arms and decrease exploratory behavior in males (Padovan et al., 2000; Padovan and Guimarães, 2000; Reis et al., 2011). However, limited information is available in the literature for females.

In the elevated plus maze (EPM) test, the percentage time spent in open arms decreased after stress in males in comparison to controls (Figure 2B; One-way ANOVA, F_2, 21_=8.510, p<0.05). Post hoc analysis shows a significant decrease between ctrl and RS rats, and between ctrl and RS24 rats. There were no changes in the number of entries in open arms in comparison to ctrl (figure 2C; One-way ANOVA, F_2, 21_=2.957; p=0.0739).

**Figure 2:**
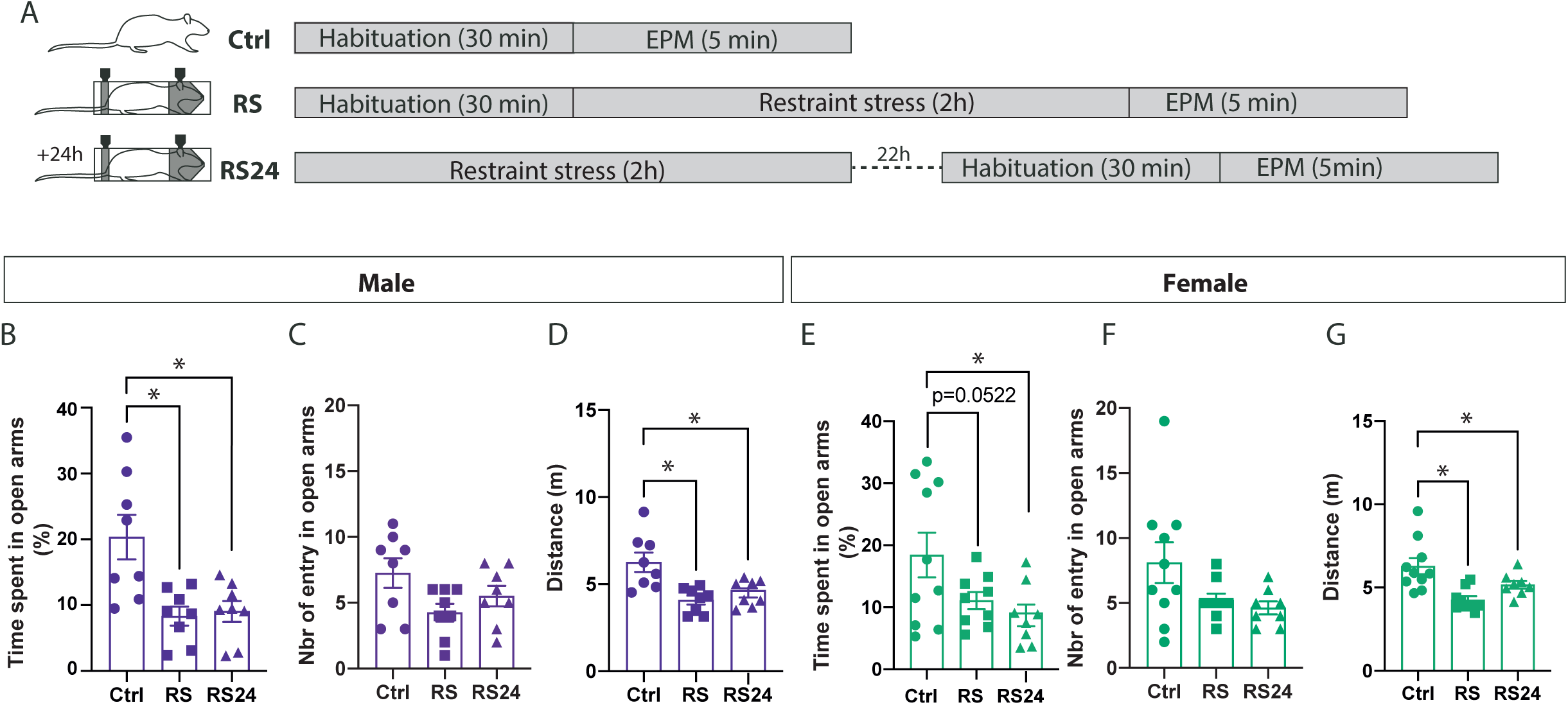
Effect of restraint stress on avoidance and exploratory behavior. **(A)** Experimental groups and procedures for behavioral evaluation in elevated plus maze (EPM) in control (Ctrl) and restraint stressed rats immediately (RS) or 24h after stress (AR24). Quantification of behaviors with percentage of time spent in open arms **(B, E)**, number of entries in open arms (C, F) and distance in meter **(D, G)** in males (left) and females (right) (mean ± SEM). Male Ctrl, n=8, Male RS, n=8, Male RS24, n=8; Female Ctrl, n=10; Female RS, n=9, Female RS24, n=8; *p<0.05.

In females, the percentage time spent in open arms decreases significantly after stress in comparison to ctrl (Figure 2D; One-way ANOVA, F_2, 24_=7.504, p<0.05). Post hoc analysis shows a trend towards a decrease between ctrl and RS rats (p=0.0522), and a significant difference between ctrl and RS24 rats (p<0.05). As in males, stress did not induce significant changes in the number of entries in open arms (figure 2F; One-way ANOVA, F_2, 24_=3:06; p=0.0655).

Exploratory behavior was investigated by measuring the distance travelled in the EPM (Figure 2D, G). In both males and females, the distance was decreased immediately and 24h after stress (One-way ANOVA; males: F_2, 21_=2.887, p<0.05; females: F_2, 24_=1.594, p<0.05). Post hoc analysis showed a difference between ctrl and RS rats and between ctrl and RS24 rats in both males and females (p<0.05). This suggests that acute stress increases avoidance in both male and female rats.

### The number of neurons projecting from the aIC to the NAcC does not differ between naive males and females

Previous studies have demonstrated that dimorphism exists between males and females for brain volume and surface area and reported that men display global larger brain volume (Jäncke et al., 2015). However, for some brain areas such as the insula, females reported larger volume on both sides of their brain (Longarzo et al., 2021). Although connectivity of the aIC has been thoroughly described in rodents (Gehrlach et al., 2020), it is not clear if there is a difference in aIC projection neurons between males and females, in particular in structures impacted by stress, such as the NAcC. Thus, we measured the percentage of CTB positive neurons (% of CTB+/NeuN+ cells) of the aIC-NAcC pathway in male and female naive rats following an injection of retrograde CTB tracer in the NAcC (figure 3A) and found no differences between male and female rats in the density of inputs in this pathway (Figure 3D, Mann-Whitney, U=8, p=0.4206, n=5 males and n=5 females).

**Figure 3:**
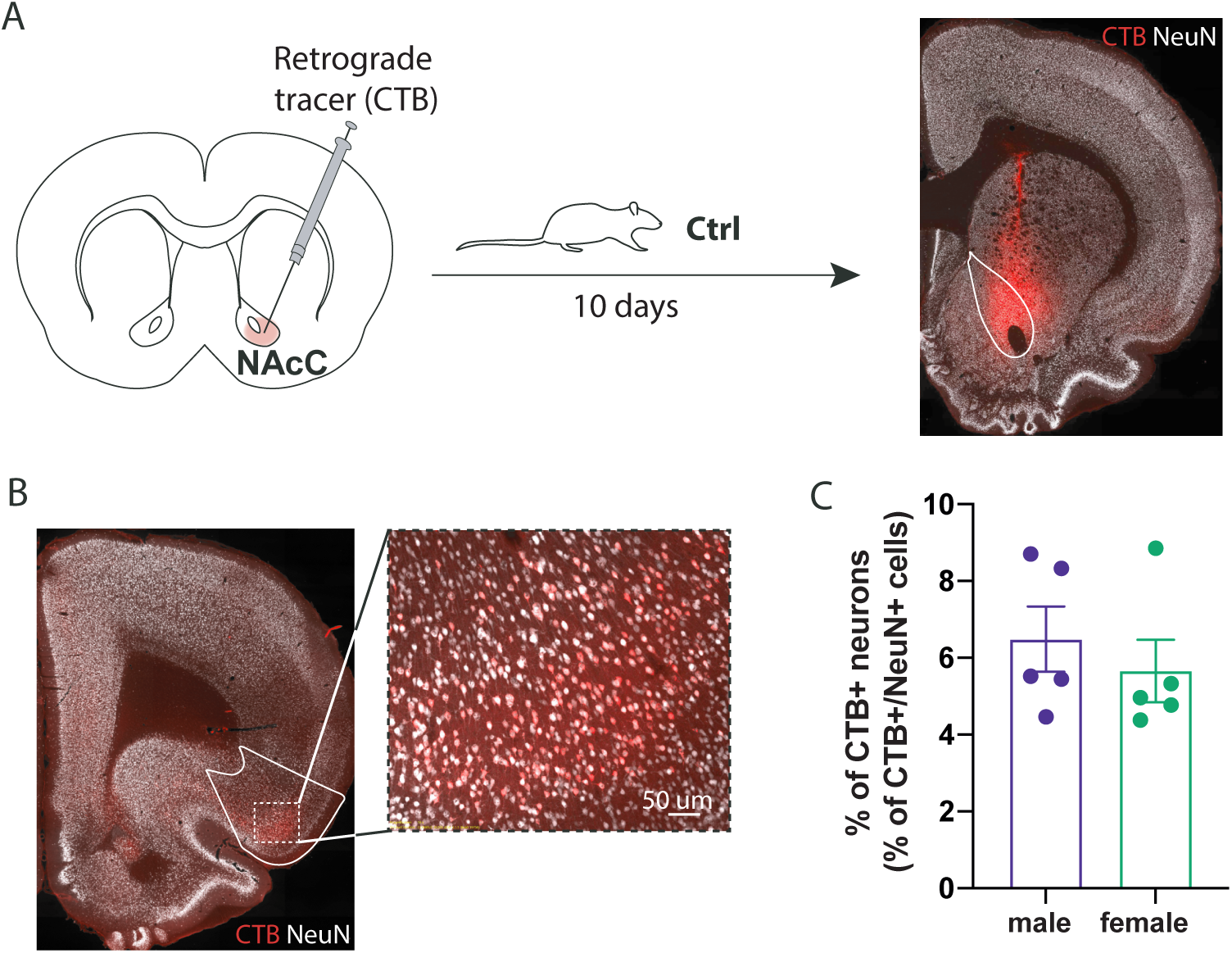
Density of inputs from the aIC to the NAcC in males and females. **(A)** Schematic representation of experimental procedure for NAcC CTB injection in male and female rats (left) and representative fluorescence microscopy image of the CTB injection site in a coronal slice of the NAcC showing CTB and NeuN labelling (right). **(B)** Representative fluorescence microscopy image of coronal slice of the aIC showing CTB and NeuN labelling **(C)** Percentage of CTB positive neurons in males and females in the aIC (mean ± SEM). Male, n=5; Female, n=5.

### Insula-accumbens synaptic plasticity is different between naive males and females

To examine synaptic plasticity in the aIC-NAcC pathway in females and males, aIC-evoked activity was assessed in the NAcC after HFS to the aIC (figure 4A).

**Figure 4:**
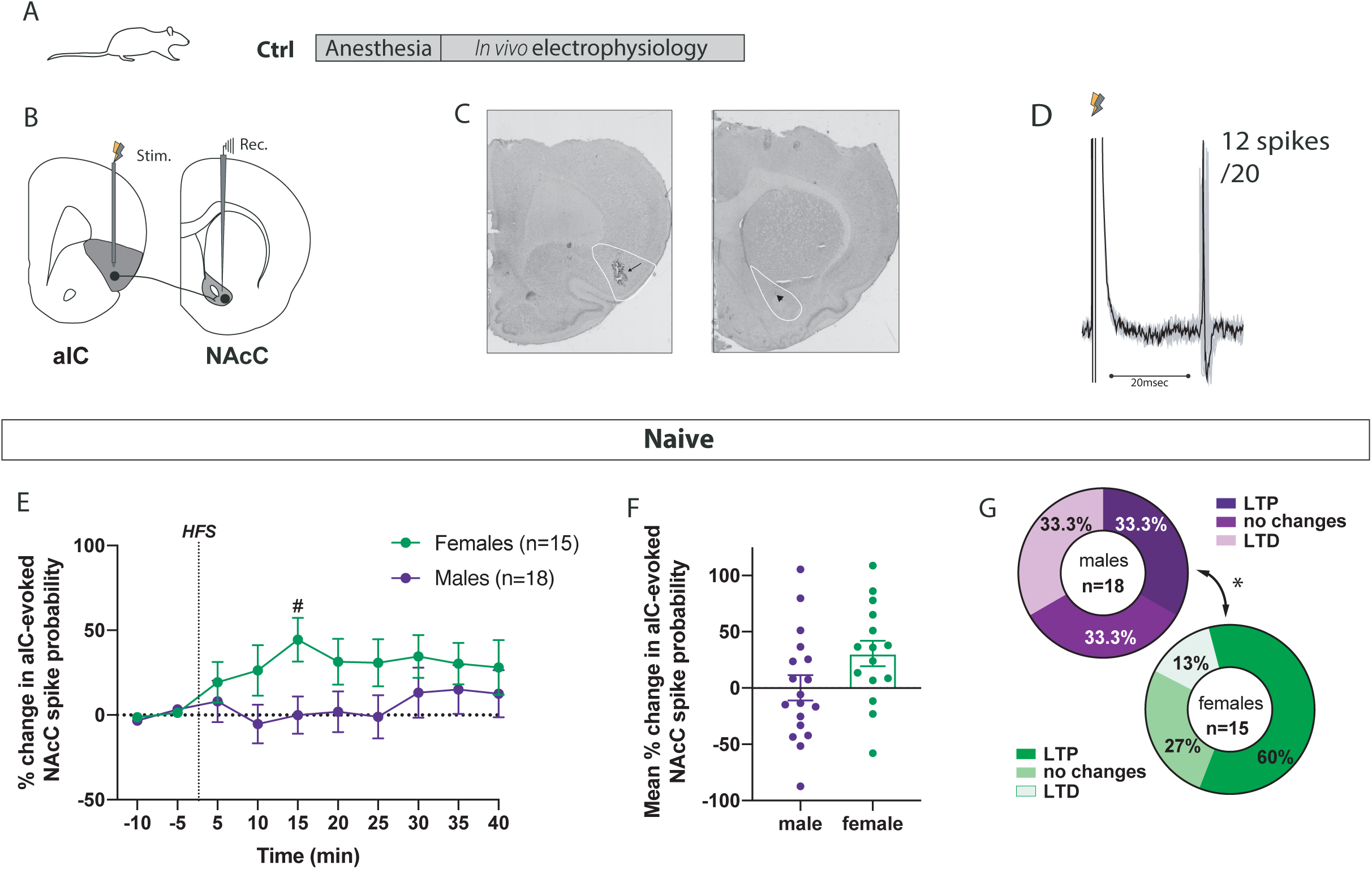
Neuronal plasticity of the anterior insular cortex to nucleus accumbens core pathway in male and female rats. **(A)** Experimental group and procedure for *in vivo* electrophysiology in anesthetized control rats (Ctrl) **(B)** Schematic of the placements of the stimulation electrode in the anterior insular cortex (aIC) and the recording electrode in the core of the nucleus accumbens (NAcC). Grey lightning represents single pulse stimulation and yellow lightning represents high-frequency stimulation (HFS). **(C)** Representative aIC stimulation (black arrow) and NAcC recording sites (black triangle). **(D)** Representative electrophysiological trace showing aIC-evoked spike probability recorded from a NAcC neurons in a Ctrl male rat with an overview of 20 frames. The lightning indicates the stimulation artifact; 12 spikes out of 20 stimulations were evoked. **(E)** Mean percentage change (± SEM) in aIC-evoked spike probability, normalized to the baseline, after HFS to the aIC in males (purple circles) and females (green circles) **(F)** Mean percent change in aIC-evoked responses (± SEM) following HFS in male (purple) and female (green) rats. (G) Proportions of recorded neurons showing an LTP, no changes or LTD in males (purple) and female rats (green). Male n=18; Female Ctrl. *p<0.05; ^#^p<0.05 in comparison to baseline.

No differences were found for the baseline current intensity (Kruskal-Wallis test, K=7.628, p=0.3665), baseline spike probability (One-Way ANOVA, F_7, 75_=1.067, p=0.3930) and mean latency (Kruskal-Wallis test, K=8.612, p=0.2817) for the neurons recorded in the NAcC of male and female rats (Table 1).

**Table 1:** Latency to spike discharge and aIC stimulus intensity and baseline spike probability of neurons recording in the NAcC during the baseline period of animals.

|  | Males |  |  |  | Females |  |  |  |
| --- | --- | --- | --- | --- | --- | --- | --- | --- |
|  | Ctrl | RS- | RS+ | RS24 | Ctrl | RS- | RS+ | RS24 |
| Number of neurons | 18 | 8 | 6 | 9 | 15 | 5 | 9 | 13 |
| Latency to spike discharge (ms) | 19.12 ± 0.90 | 22.11 ± 1.17 | 20.84 ± 2.37 | 22.19 ± 1.26 | 20.68 ± 1.34 | 22.11 ± 1.27 | 23.10 ± 1.47 | 19.70 ± 0.84 |
| Current intensity (mA) | 0.95 ± 0.06 | 1.02 ± 0.12 | 0.95 ± 0.12 | 0.98 ± 0.1 | 0.96 ± 0.06 | 0.66 ± 0.11 | 0.97 ± 0.09 | 0.86 ± 0.09 |
| Baseline spike probability | 0.60 ± 0.04 | 0.53 ± 0.07 | 0.44 ± 0.05 | 0.54 ± 0.06 | 0.48 ± 0.04 | 0.53 ± 0.03 | 0.51 ± 0.05 | 0.50 ± 0.05 |

HFS applied to the aIC had a different effect on aIC-induced spike probability in control female and male rats (Figure 4E, F). Indeed, a mixed effect analysis indicated an interaction between time and sex (F_9, 258_=2.233, p<0.05), although no effect of time (F_9, 258_=2.233, p=0.1240) and no effect of sex (F_1, 31_=3.221, p=0.085). Post hoc analysis showed a significant increase of spike probability 15 minutes post-HFS in comparison to baseline in females, but no changes in males (Figure 4E).

A strong inter-individual variability was observed in the ctrl groups for males and females. We thus clustered the different response types in relation to the mean percentage change in spike probability after HFS. In males, aIC HFS induced LTP in 33% of recorded neurons, LTD in 33 % and no changes in 33 %. In females, 60% of recorded neurons showed an LTP post-HFS, 13.3% an LTD, and 26.7 % no changes. The proportion of responses was different between males and females (X^2^, p<0.05) (figure 4G). It should be noted that this high variability did not depend on the location of recording and/or stimulating electrode (supplementary figure 2).

This suggests that under control conditions, high frequency stimulation of the aIC is more efficient in males than in females to induce long-term depression and is more efficient in females than in males to induce long-term potentiation.

### Restraint stress induces lasting changes in synaptic plasticity in the aIC-NAcC pathway in male, but not female, rats

To examine potential changes of synaptic transmission in the aIC-NAcC pathway in stressed females and males, aIC-evoked activity was assessed in the NAcC after HFS to the aIC in Ctrl, RS and RS24 rats (figure 5).

**Figure 5:**
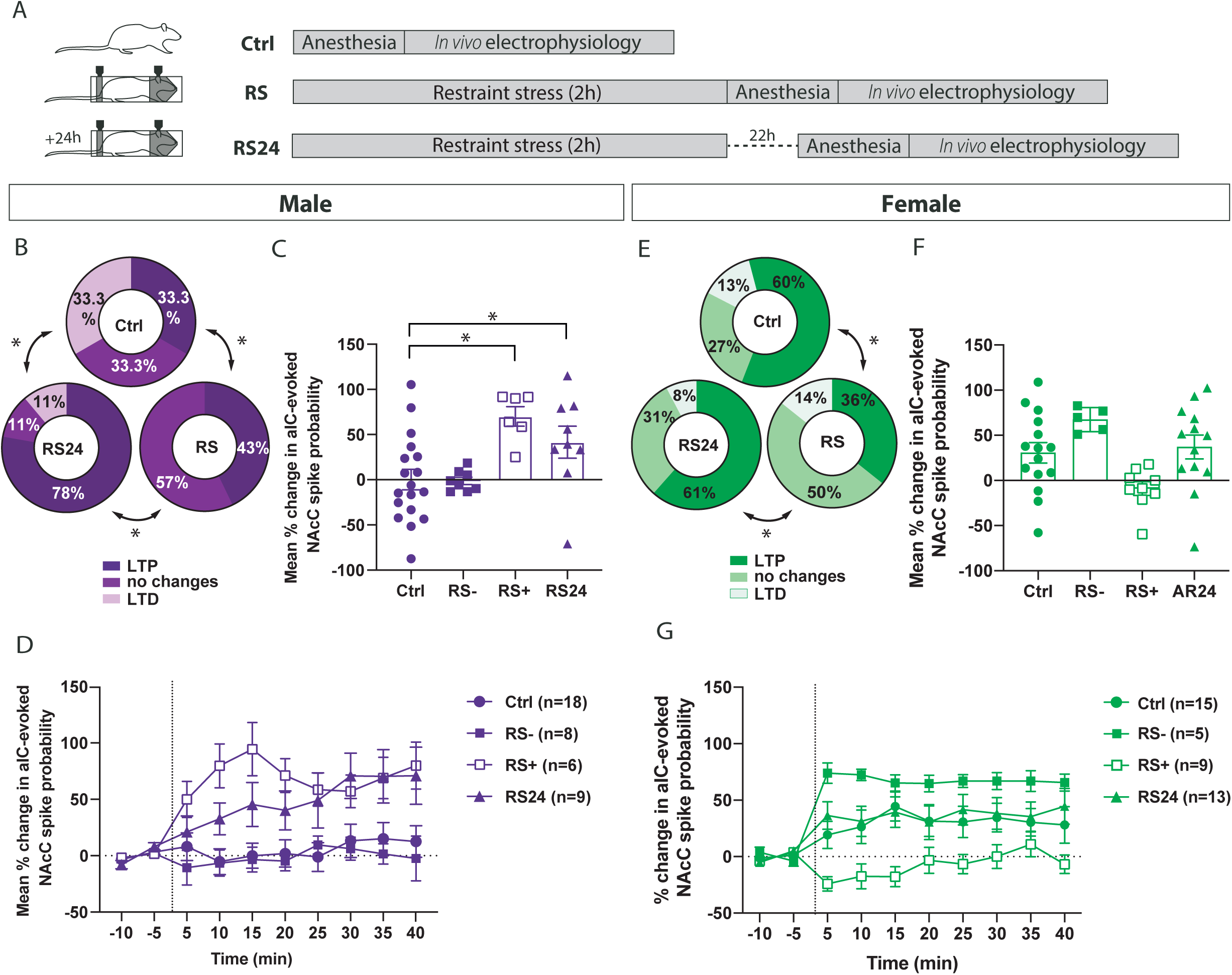
Effect of restraint stress on neuronal plasticity of the anterior insular cortex to nucleus accumbens core pathway in male and female rats. **(A)** Experimental groups and procedures for *in vivo* electrophysiology in anesthetized control rats (Ctrl) and restraint stressed rats anesthetized immediately (RS) or 24h after stress (RS24). **(B, E)** Proportions of recorded neurons showing an LTP, no changes or LTD in Ctrl, after RS and 24h after RS (RS24) males (purple, B) and female rats (green, E). (C) Mean percentage change (± SEM) in aIC-evoked spike probability following HFS to the aIC in ctrl (circle), immediately after stress, RS-(no changes), immediately after stress, RS+ (LTP) and 24h after stress (RS24) **(F)** Mean percent change in aIC-evoked responses (± SEM) following HFS in ctrl (circle) female (green) rats, immediately (RS, square) and 24h after restraint stress (RS24, triangle). **(D, G)** Mean percentage change (± SEM) in aIC-evoked spike probability, normalized to the baseline, after HFS to the aIC) in Ctrl (circle), RS-(full square), RS+ (open square) and RS24 (triangle) in male (purple, D) and female (green, G) rats. Male Ctrl, n=18; Male RS, n= 14, Male RS24, n=9; Female Ctrl, n=15, Female RS, n=14, Female RS24, n=13. *p<0.05.

Considering the inter-individual variability observed in the ctrl groups, we clustered the different response types in relation to the mean percentage change in spike probability after HFS in the restraint stress groups.

In males, immediately after restraint stress, aIC HFS induced LTP in 43 % of recorded neurons and no changes in 57 %. Twenty-four hours after RS, aIC HFS induced LTP in 78 % of recorded neurons, and 11 % of LTD or no changes (Figure 5B). The proportion of responses was significantly different between control and RS males, control and RS24 males and RS and RS24 males (X^2^; p<0.05) (figure 5B). We thus divided the two types of responses in the RS group into RS-(no changes) and RS+ (LTP). One-way ANOVA analysis of the mean percentage change across the 40 min period post-HFS showed a significant difference in spike probability (p<0.05). Post hoc analyses showed that immediately after stress, there was a significant difference between ctrl and RS+ and a trend toward difference (p=0.06) between ctrl and RS 24 (Figure 5C). A mixed effect analysis indicated an effect of time (F_27, 307_=6.901, p<0.05), an effect of condition (naive or stressed; F_3, 37_=6.325, p<0.05) and interaction between condition and time (F_27, 31307_=2.739, p<0.05) (Figure 5D).

In female rats, immediately after restraint stress, aIC HFS induced LTP in 36 % of recorded neurons, no changes in 50% and an LTD in 14 %. Twenty-four hours after RS, aIC HFS induced LTP in 61% of recorded neurons, and LTD in 8 % and no changes in 31 % (Figure 5E). The proportion of responses was significantly different between control and RS rats, and between RS and RS24 rats (X^2^; p<0.05). We applied the same criterion used for males to divide the types of responses in the RS group into RS-(LTP, >20% change and RS+ (<20%change, no changes). A One-Way ANOVA analysis of the mean percentage change across the 40 min period showed no change in post-HFS spike probability (F_2, 39_=0.6221, p=0.5421) (Figure 5F). A One-Way ANOVA analysis of the mean percentage change across the post-HFS 40 min period showed a significant difference in spike probability (p<0.05). Post hoc analyses showed that immediately after stress, there was a trend toward a difference between ctrl and RS+ (p=0.06) (Figure 5F). A mixed effect analysis indicated an effect of time (F_27, 329_=10.56, p<0.05), an effect of condition (naive or stressed; F_3, 38_=4.366, p<0.05) and an interaction between condition and time (F_27, 329_=2.712, p<0.05) (Figure 5G). Altogether these results suggest that in males, stress induces a loss of synaptic mechanisms that allows long-term depression of excitability, which lasts up to 24h. These results also suggest that in females, stress induces a change in the proportion of neurons displaying LTP after restraint, which does not last.

## DISCUSSION

To our knowledge, this is the first study to examine synaptic plasticity in the insular cortex-nucleus accumbens pathway in male and female rats and the impact of stress on aIC information processing in the NAc.

While most studies on stress have been conducted in males, less is known about the neurobiological effects of stress in females. In this study, we used a two-hour immobilization stress in male and female rats and we found an increase in cFos expression in the PVN after acute restraint stress, consistent with previous studies (Kovács et al., 2018; Mohammad et al., 2000).

In the EPM, we found a decrease in the % time spent in open arms, as well as a decrease in exploratory behavior in males and females. However, we found no changes in the number of entries in open arms. It should be noted that female rat behavior in the EPM has been shown to be characterized primarily by activity, whereas male behavior has been characterized primarily by anxiety (Fernandes et al., 1999). Our data suggests that in both males and females, restraint stress reduces activity, as well as reduces the time spent in anxiogenic places, such as open arms that can last up to 24h, which is consistent with previous studies showing a lasting effect of stress in males (Mitra et al., 2005). This enhanced avoidance, and decreased exploratory behaviorcan be described as hypervigilance in anticipation of a threat, triggered by acute stress (Endler and Kocovski, 2001).

Our electrophysiological data indicate different information processing in the NAc from aIC inputs between naive males and females. In naive rats, HFS of the aIC induced no overall changes in spike probability in males but an overall long-term potentiation in females. This is due in part to a large proportion of neurons recorded in males that display an LTD (33 %), whereas only a small proportion of neurons recorded in females showed this type of plasticity. This suggests that under basal, non-stressed conditions, the aIC-NAc pathway in males can be potentiated or depressed, whereas in females, the mechanisms that allow depression seem lacking. Therefore, information from the aIC, such as emotional awareness (Gu et al., 2013), would be strengthened in females in basal conditions, increasing information processing in the NAcC. This would be consistent with previous studies demonstrating that females have greater emotional awareness than males (Barrett et al., 2000).

This difference in synaptic plasticity between naive males and females could not be due to a difference in the number of neurons projecting from the aIC to the NAcC, since the number of CTb-positive neurons in the aIC is the same between naive males and females. However, it is possible that the density of aIC synaptic terminals is different in the NAcC between males and females, as described in other structures (Carvalho-Netto et al., 2011). Indeed, an increased number of synaptic inputs, which increases the probability to induce potentiation (Nicoll et al., 1988), has been shown in females in comparison to males in some structures (Carvalho-Netto et al., 2011). Therefore, further analysis would be necessary to determine presynaptic nerve terminal density in the NAcC of males and females. Another possibility would be sex differences in NAcC synaptic connectivity, in particular in dendritic spine density. Indeed, neuroanatomical sex differences have been shown in the NAcC. Greater spine density and greater head spine size have been described in females, in comparison to males, providing evidence of differences in structural synaptic connectivity of glutamatergic inputs in this region (Forlano and Woolley, 2009). This sex difference might indicate difference in synaptic plasticity in basal conditions. It has also been shown that females had a higher proportion of spines with large heads near tyrosine-hydroxylase immunoreactive fibers suggesting a potential sex differences in dopaminergic modulation of large spines synapses (Wissman et al., 2012). Since dopamine is a potent modulator of synaptic plasticity in the NAcC (Goto and Grace, 2005), different dopaminergic modulation in the NAcC between males and females could potentially induce different synaptic plasticity in the NAcC in basal conditions.

Another possible explanation is the difference in the endocannabinoid (eCB) system between males and females. Indeed, levels of eCB (andamide and 2-arachidonoylglycerol, 2-AG) are different in males and females in some brain areas such as the amygdala (Blanton et al., 2021), as well as affinity for CB1 receptors are different in males and females in the striatum (de Fonseca et al., 1994). Considering that eCB-dependent plasticity has been shown to be induced by tetanus stimulation in the NAc (Robbe et al., 2002), it is possible that our protocol of stimulation induces eCB-dependent LTD in males and eCB-dependent LTP in females (Piette et al., 2020), and that differences in eCB or eCB receptors in the accumbens of males and females induces differences in synaptic plasticity between males and females in the aIC-NAcC pathway.

Whereas restraint stress impacted for at least 24h the aIC-NAcC pathway in males, it had a transient effect in females. Indeed, in males, we demonstrated a total loss of LTD after RS, revealing two types of responses, a potentiation of aIC-NAcC synaptic efficacy and no changes, which last at least 24h. In females, the proportions of synaptic plasticity after stress were different, with less neurons displaying LTP than control animals. Previous studies have shown sex-specific role for corticotropin-releasing factor (CRF), released by PVN neurons in response to a stressor or a stressed conspecific, as a modulator of aIC synaptic activity (Rieger et al., 2022). Indeed, CRF injected in the aIC induced a depolarization of pyramidal neurons in both males and females and a reduction in presynaptic inhibitory tone in males only. Therefore, in our study, CRF release in response to restraint stress could affect the aIC-NAc pathway in a different manner in males and females, by suppressing LTD mechanisms in males and decreasing LTP mechanisms in females.

Previous studies have shown that spike-timing dependent eCB-dependent LTD in the NAc observed in naive animals is attenuated after stress in anxious animals (Bosch-Bouju et al., 2016), which was restored by increasing availability of 2-AG. Although in our study we did not directly investigate role of eCB and the electrophysiological measures were different between studies, our data support the idea that stress enhances avoidance and blocks LTD in the aIC-NAc pathway, possibly via the eCB system, only in males. Depending on CB receptor expression and eCB levels, eCB-LTP can be induced heterosynaptically or homosynaptically (Piette et al., 2020). It is therefore possible that in the same pathway, eCB plasticity differs between males and females, which would be impacted differently by stress. This will need to be further investigated.

## CONCLUSION

Taken together, our results suggest that stress increases information processing from the aIC to the NAcC only in males, which could be a ‘buffer’ coping mechanism in a stressful situation (Lea et al., 2019). A decrease of potentiation by acute stress in females would suggest that emotional awareness, already present and processed in basal conditions, could not be further used as a coping mechanism.

This work provides new insights into the neuroadaptations in the aIC-NAcC network induced by stress in female and male rats.

## Acknowledgements

This work has benefited from the facilities and expertise of PREBIOS platform (University of Poitiers). This work was supported by the Institute National de la Santé et de la Recherche Médicale, the Centre National pour la Recherche Scientifique, the University of Poitiers, CHU of Poitiers, and the Région Nouvelle-Aquitaine. The authors thank Catherine Le Goff for technical support.

## Notes

### Competing Interest Statement

The authors have declared no competing interest.

## References

Bangasser, D.A., Valentino, R.J., 2014. Sex differences in stress-related psychiatric disorders: Neurobiological perspectives. Frontiers in Neuroendocrinology, Sex Differences in Neurological and Psychiatric Disorders 35, 303–319. 10.1016/j.yfrne.2014.03.008

Bangasser, D.A., Wiersielis, K.R., 2018. Sex differences in stress responses: a critical role for corticotropin-releasing factor. Hormones (Athens) 17, 5–13. 10.1007/s42000-018-0002-z

Bankhead, P., Loughrey, M.B., Fernández, J.A., Dombrowski, Y., McArt, D.G., Dunne, P.D., McQuaid, S., Gray, R.T., Murray, L.J., Coleman, H.G., James, J.A., Salto-Tellez, M., Hamilton, P.W., 2017. QuPath: Open source software for digital pathology image analysis. Sci Rep 7, 16878. 10.1038/s41598-017-17204-5

Barrett, L.F., Lane, R.D., Sechrest, L., Schwartz, G.E., 2000. Sex differences in emotional awareness. Personality and Social Psychology Bulletin 26, 1027–1035. 10.1177/01461672002611001

Blanton, H.L., Barnes, R.C., McHann, M.C., Bilbrey, J.A., Wilkerson, J.L., Guindon, J., 2021. Sex differences and the endocannabinoid system in pain. Pharmacol Biochem Behav 202, 173107. 10.1016/j.pbb.2021.173107

Bosch-Bouju, C., Larrieu, T., Linders, L., Manzoni, O.J., Layé, S., 2016. Endocannabinoid-Mediated Plasticity in Nucleus Accumbens Controls Vulnerability to Anxiety after Social Defeat Stress. Cell Reports 16, 1237–1242. 10.1016/j.celrep.2016.06.082

Carvalho-Netto, E.F., Myers, B., Jones, K., Solomon, M.B., Herman, J.P., 2011. Sex differences in synaptic plasticity in stress-responsive brain regions following chronic variable stress. Physiol Behav 104, 242–247. 10.1016/j.physbeh.2011.01.024

Chajut, E., Algom, D., 2003. Selective attention improves under stress: implications for theories of social cognition. J Pers Soc Psychol 85, 231–248. 10.1037/0022-3514.85.2.231

Chrousos, G.P., Gold, P.W., 1992. The concepts of stress and stress system disorders. Overview of physical and behavioral homeostasis. JAMA 267, 1244–1252.

Craig, A.D., 2009. How do you feel--now? The anterior insula and human awareness. Nature reviews. Neuroscience 10, 59–70. 10.1038/nrn2555

Craig, A.D., 2002. How do you feel? Interoception: the sense of the physiological condition of the body. Nature reviews. Neuroscience 3, 655–66. 10.1038/nrn894

Critchley, H.D., Wiens, S., Rotshtein, P., Ohman, A., Dolan, R.J., 2004. Neural systems supporting interoceptive awareness. Nature neuroscience 7, 189–95. 10.1038/nn1176

de Fonseca, F.R., Cebeira, M., Ramos, J.A., Martín, M., Fernández-Ruiz, J.J., 1994. Cannabinoid receptors in rat brain areas: Sexual differences, fluctuations during estrous cycle and changes after gonadectomy and sex steroid replacement. Life Sciences 54, 159–170. 10.1016/0024-3205(94)00585-0

Endler, N.S., Kocovski, N.L., 2001. State and trait anxiety revisited. J Anxiety Disord 15, 231–245. 10.1016/s0887-6185(01)00060-3

Fernandes, C., González, M.I., Wilson, C.A., File, S.E., 1999. Factor Analysis Shows That Female Rat Behaviour Is Characterized Primarily by Activity, Male Rats Are Driven by Sex and Anxiety. Pharmacology Biochemistry and Behavior 64, 731–736. 10.1016/S0091-3057(99)00139-2

Floresco, S.B., 2015. The nucleus accumbens: an interface between cognition, emotion, and action. Annu Rev Psychol 66, 25–52. 10.1146/annurev-psych-010213-115159

Floresco, S.B., Blaha, C.D., Yang, C.R., Phillips, A.G., 2001. Modulation of hippocampal and amygdalar-evoked activity of nucleus accumbens neurons by dopamine: cellular mechanisms of input selection. J Neurosci 21, 2851–60.

Forlano, P., Woolley, C., 2009. Quantitative analysis of pre- and postsynaptic sex differences in the nucleus accumbens - Forlano - 2010 - Journal of Comparative Neurology - Wiley Online Library [WWW Document]. URL https://onlinelibrary.wiley.com/doi/10.1002/cne.22279 (accessed 4.26.23).

Gehrlach, D.A., Gaitanos, T.N., Klein, A.S., Weiand, C., Hennrich, A.A., Conzelmann, K.-K., Gogolla, N., 2020. A whole-brain connectivity map of mouse insular cortex. bioRxiv 2020.02.10.941518. 10.1101/2020.02.10.941518

Gill, K.M., Grace, A.A., 2013. Differential effects of acute and repeated stress on hippocampus and amygdala inputs to the nucleus accumbens shell. Int J Neuropsychopharmacol 1–13.

Goto, Yukiori, Grace, A.A., 2005. Dopamine-Dependent Interactions between Limbic and Prefrontal Cortical Plasticity in the Nucleus Accumbens: Disruption by Cocaine Sensitization. Neuron 47, 255–266. 10.1016/j.neuron.2005.06.017

Goto, Y., Grace, A.A., 2005. Dopaminergic modulation of limbic and cortical drive of nucleus accumbens in goal-directed behavior. Nature neuroscience 8, 805–12.

Gu, X., Hof, P.R., Friston, K.J., Fan, J., 2013. Anterior Insular Cortex and Emotional Awareness. J Comp Neurol 521, 3371–3388. 10.1002/cne.23368

Jäncke, L., Mérillat, S., Liem, F., Hänggi, J., 2015. Brain size, sex, and the aging brain. Hum Brain Mapp 36, 150–169. 10.1002/hbm.22619

Khoury, N.M., Lutz, J., Schuman-Olivier, Z., 2018. Interoception in Psychiatric Disorders: A Review of Randomized Controlled Trials with Interoception-based Interventions. Harv Rev Psychiatry 26, 250–263. 10.1097/HRP.0000000000000170

Kovács, L.Á., Schiessl, J.A., Nafz, A.E., Csernus, V., Gaszner, B., 2018. Both Basal and Acute Restraint Stress-Induced c-Fos Expression Is Influenced by Age in the Extended Amygdala and Brainstem Stress Centers in Male Rats. Front Aging Neurosci 10, 248. 10.3389/fnagi.2018.00248

Lea, R.G., Davis, S.K., Mahoney, B., Qualter, P., 2019. Does Emotional Intelligence Buffer the Effects of Acute Stress? A Systematic Review. Frontiers in Psychology 10.

Longarzo, M., Mele, G., Alfano, V., Salvatore, M., Cavaliere, C., 2021. Gender Brain Structural Differences and Interoception. Front Neurosci 14, 586860. 10.3389/fnins.2020.586860

Mitra, R., Jadhav, S., McEwen, B.S., Vyas, A., Chattarji, S., 2005. Stress duration modulates the spatiotemporal patterns of spine formation in the basolateral amygdala. Proceedings of the National Academy of Sciences of the United States of America 102, 9371–6. 10.1073/pnas.0504011102

Mogenson, G.J., Jones, D.L., Yim, C.Y., 1980. From motivation to action: Functional interface between the limbic system and the motor system. Progress in Neurobiology 14, 69–97. 10.1016/0301-0082(80)90018-0

Mohammad, G., Chowdhury, I., Fujioka, T., Nakamura, S., 2000. Induction and adaptation of Fos expression in the rat brain by two types of acute restraint stress. Brain Research Bulletin 52, 171–182. 10.1016/S0361-9230(00)00231-8

Naqvi, N.H., Gaznick, N., Tranel, D., Bechara, A., 2014. The insula: a critical neural substrate for craving and drug seeking under conflict and risk. Ann N Y Acad Sci 1316, 53–70. 10.1111/nyas.12415

Nicoll, R.A., Kauer, J.A., Malenka, R.C., 1988. The current excitement in long-term potentiation. Neuron 1, 97–103. 10.1016/0896-6273(88)90193-6

Paxinos, G., Watson, C., 2017. The Rat Brain in Stereotatic Coordinates. Academic Press, San Diego.

Piette, C., Cui, Y., Gervasi, N., Venance, L., 2020. Lights on Endocannabinoid-Mediated Synaptic Potentiation. Frontiers in Molecular Neuroscience 13.

Prentice, F., Hobson, H., Spooner, R., Murphy, J., 2022. Gender differences in interoceptive accuracy and emotional ability: An explanation for incompatible findings. Neuroscience & Biobehavioral Reviews 141, 104808. 10.1016/j.neubiorev.2022.104808

Rieger, N.S., Varela, J.A., Ng, A.J., Granata, L., Djerdjaj, A., Brenhouse, H.C., Christianson, J.P., 2022. Insular cortex corticotropin-releasing factor integrates stress signaling with social affective behavior. Neuropsychopharmacology 47, 1156–1168. 10.1038/s41386-022-01292-7

Robbe, D., Kopf, M., Remaury, A., Bockaert, J., Manzoni, O.J., 2002. Endogenous cannabinoids mediate long-term synaptic depression in the nucleus accumbens. Proc Natl Acad Sci U S A 99, 8384–8388. 10.1073/pnas.122149199

Schneider, C.A., Rasband, W.S., Eliceiri, K.W., 2012. NIH Image to ImageJ: 25 years of image analysis. Nat Methods 9, 671–675. 10.1038/nmeth.2089

Sliz, D., Hayley, S., 2012. Major Depressive Disorder and Alterations in Insular Cortical Activity: A Review of Current Functional Magnetic Imaging Research. Front Hum Neurosci 6, 323. 10.3389/fnhum.2012.00323

Smith, K.E., Pollak, S.D., 2020. Early life stress and development: potential mechanisms for adverse outcomes. J Neurodev Disord 12, 34. 10.1186/s11689-020-09337-y

Stevenson, C.W., Gratton, A., 2003. Basolateral amygdala modulation of the nucleus accumbens dopamine response to stress: role of the medial prefrontal cortex. Eur J Neurosci 17, 1287–1295. 10.1046/j.1460-9568.2003.02560.x

Terasawa, Y., Shibata, M., Moriguchi, Y., Umeda, S., 2013. Anterior insular cortex mediates bodily sensibility and social anxiety. Soc Cogn Affect Neurosci 8, 259–266. 10.1093/scan/nss108

Thompson, A.E., Voyer, D., 2014. Sex differences in the ability to recognise non-verbal displays of emotion: A meta-analysis. Cognition and Emotion 28, 1164–1195. 10.1080/02699931.2013.875889

Wissman, A.M., May, R.M., Woolley, C.S., 2012. Ultrastructural analysis of sex differences in nucleus accumbens synaptic connectivity. Brain Struct Funct 217, 181–190. 10.1007/s00429-011-0353-6

