## supplemental Figure 1 for "Sex-dependent effects of stress on insular cortex-to-nucleus accumbens synaptic plasticity"

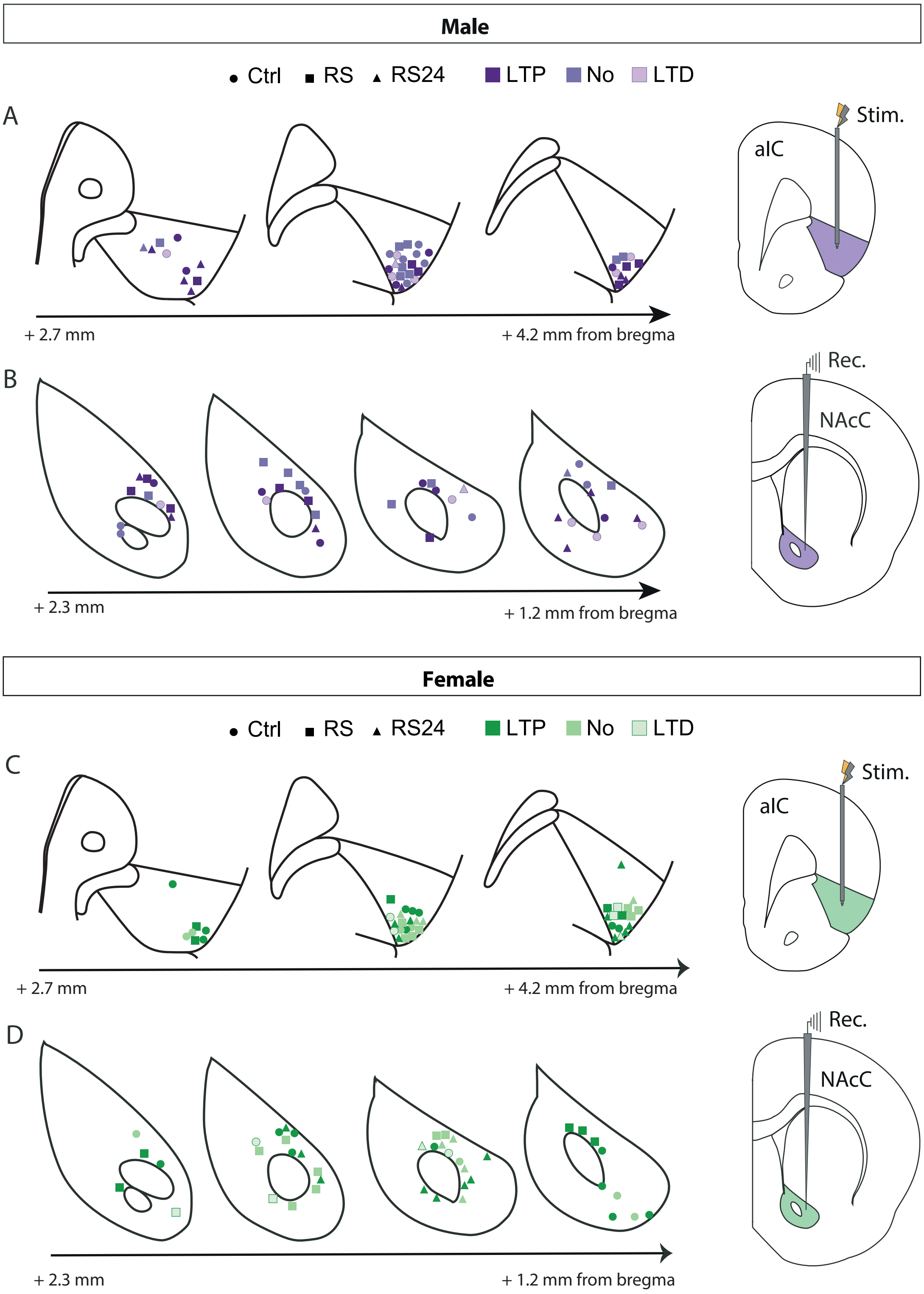


**Supplementary Figure 1:** Placements of stimulation (A, C) in the anterior insular cortex (aIC) and recording in the nucleus accumbens core (NAcC) (B, D) electrodes in males (purple) and females (green). Colours are accorded to plasticity response type (LTP, absence of plasticity or LTD).
