## supplemental Figure 2 for "Sex-dependent effects of stress on insular cortex-to-nucleus accumbens synaptic plasticity"

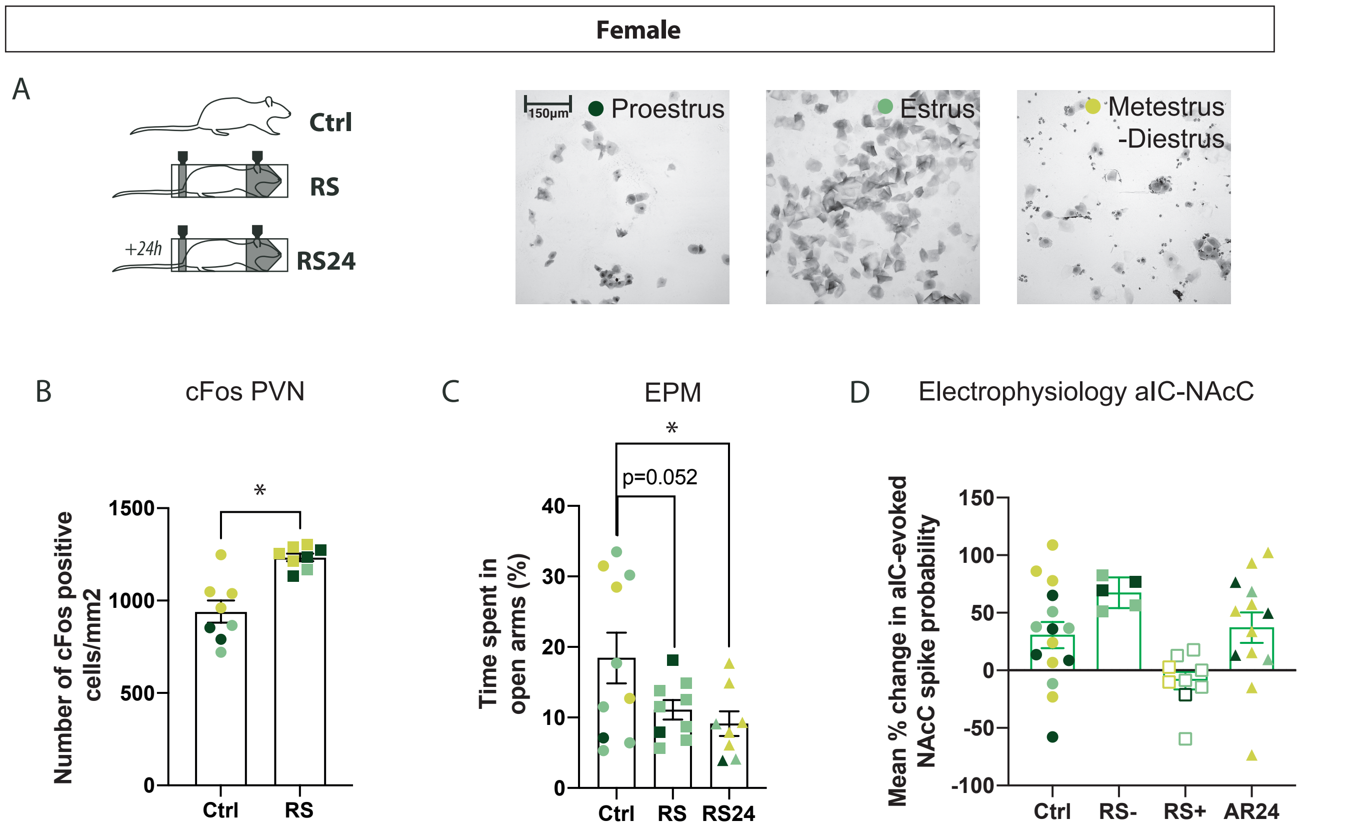


Supplementary Figure 2: Identification of estrous stages of females for all experiments.
(A) Experimental groups and representative image of cytological sample of vaginal smears to identify estrous stage (proestrus, estrus and metestrus/diestrus). Impact of restraint stress on the mean +/- SEM cFos density in the PVN (B) and on the percentage of time spend in open arms in the EPM (C).
(D) Quantification of mean percentage +/- SEM of aIC- evoked spike probability, normalized to baseline, after HFS in the aIC. *p<0.05
